# Executive dysfunction is associated with altered hippocampal-prefrontal functional connectivity in male 3xTg Alzheimer’s model mice

**DOI:** 10.1101/2024.09.26.615115

**Authors:** Grace Cunliffe, Li Yang Tan, Jung Sangyong, Jonathan Turner, John Gigg

## Abstract

Executive function depends on connectivity between the ventral hippocampus and medial prefrontal cortex (mPFC). How abnormalities in this pathway lead to cognitive dysfunction in Alzheimer’s disease (AD) have yet to be elucidated. Here, male 3xTg AD mice at 6-months displayed maladaptive decision-making in the rodent 4-Choice Gambling Task measure of executive function. Extracellular field recordings in the infralimbic cortex at this age showed layer-specific reductions in response amplitude and paired-pulse ratio following activation of hippocampal input fibres, indicating changes to short-term hippocampal-prefrontal synaptic plasticity. Bulk RNA sequencing of the mPFC in 6-month-old mice identified differential gene expression changes associated with calcium ion transport, glutamatergic, GABAergic, and dopaminergic neurotransmission. Seven of these genes (*Gpm6b, Slc38a5, Ccr5, Kcnj10, Ddah1, Gad1, Slc17a8*) were also differentially expressed in 3-month mice. These results reveal a pre-clinical deficit in executive function correlating with synaptic plasticity and gene expression changes in the mPFC of male 3xTg mice.

## Introduction

Ventral hippocampal (vHIP) inputs to the medial prefrontal cortex (mPFC) are essential for controlling executive components of cognition including attention, decision-making, risk-taking behaviours, and behavioural flexibility (1–7). These inputs are glutamatergic, arising in the CA1 and subiculum regions of the vHIP and terminating predominantly in layers II/III and V of the infralimbic (IL) and prelimbic (PL) cortices of the mPFC (8–11). Both the prefrontal cortex and hippocampus display substantial levels of neurodegeneration and amyloid beta (Aβ) plaque and tau neurofibrillary tangle (NFT) accumulation over the course of Alzheimer’s disease (AD), so it is unsurprising that alongside declarative memory loss, cognitive deficits underlying executive dysfunction are amongst the most frequently reported symptoms associated with disease progression (3,12–15). AD is the most common form of dementia, currently affecting over 55 million people worldwide, and is estimated to be prevalent in 5-8% of over 60s (16–18). Nevertheless, no cure exists and current treatment options act simply to slow the spread of disease. Studies have shown that hippocampal-prefrontal connectivity is weakened in AD rodent models (19), but changes in functional connectivity between the vHIP and mPFC and how these may lead to executive dysfunction over the course of the disease have yet to be fully elucidated.

The examination of executive function in an AD context requires a test that incorporates aspects of decision-making, such as behavioural flexibility, risk-taking tendencies, and planning for long-term gain, whilst minimising handling and related stress to the animal. Accordingly, the rodent touchscreen-operant 4-Choice Gambling Task (4CGT) was chosen for this study. This task is based on the clinically validated Iowa Gambling Task (IGT), which examines executive function in human patients. Patients with damage to the dorsolateral prefrontal cortex, analogous to the mPFC of rodents, fail to achieve optimal decision-making in the IGT (20,21), as do AD patients (22–24). Both the 4CGT and IGT employ the concept of optimal choice preference for low-risk, low penalty options that maximise reward over time, versus high-risk, high-reward choice preference, which overall leads to longer punishment duration and reduced reward availability (25). The 4CGT consists of touchscreen trials in which four lit square stimuli are presented, each associated with variable volumes of reinforcement (milkshake delivery) and punishment duration (luminance inversion). This test has been deployed effectively many times in rat models (25–28) and a small number of studies have shown that mice are also capable of learning and performing the task (29–32). To our knowledge, no preclinical studies have applied the 4CGT in an AD mouse model. The triple transgenic (3xTg) mouse (33) is a widely used model for AD as it successfully recapitulates both Aβ plaque and tau NFT pathology; the two major hallmarks of the disease. Synaptic alterations and cognitive deficits have been reported to occur in 3xTg mice as early as 4 months (33,34) due to the presence of intraneuronal amyloid. Therefore, an advantage of this model is that it provides a lengthy prodromal period over which brain and behaviour changes can be measured that relate to early AD in patients. This period represents the most likely target for disease-rectifying interventions.

Studies in AD mouse models and AD patients have reported modifications to the synaptic excitatory/inhibitory (E-I) balance in the prefrontal cortex due to alterations in glutamatergic and GABAergic receptor expression, metabolism and transport (35–40). These observations occur early in disease progression, resulting in instability of the neural network, which may potentiate disease pathogenesis and resultant cognitive disturbances (41–43). One popular metric for measuring the synaptic E-I balance is the paired pulse ratio (PPR), which is commonly used to assess short-term synaptic plasticity dependent on the probability of presynaptic vesicular transmitter release. This process is governed by presynaptic calcium dynamics and vesicle release machinery underlying vesicle exocytosis, requiring the recruitment of numerous membrane-associated proteins including synaptotagmins, complexins and SNARE proteins (44). Alterations to calcium dynamics and the expression of proteins associated with vesicle exocytosis are reported to contribute towards synaptic deficits in the hippocampus and frontal regions of AD patients (45–51). However, how AD pathology impacts short-term synaptic plasticity and local field potential (LFP) generation in the vHIP-mPFC pathway specifically is not well-defined.

To uncover alterations to hippocampal-prefrontal communication underlying impaired executive performance of 6-month-old 3xTgs on the 4CGT, LFPs were recorded from infralimbic layers II/III and V in brain slices following single or paired-pulse electrical stimulation of hippocampal input fibres. This allowed the study of input-output (IO) responses (functional connectivity) and short-term synaptic plasticity, respectively. Response amplitudes to hippocampal input stimulation were significantly reduced in IL layer II/III, whilst the paired pulse ratio was significantly reduced in layer V. RNA sequencing analysis of the mPFC in 3– and 6-month-old 3xTg and control mice was also performed, to establish whether functional changes in connectivity correlated with significant gene expression changes associated with glutamatergic and GABAergic synaptic transmission, dopaminergic neuromodulation, and calcium ion dynamics. This allowed the identification of candidates for preclinical drivers of pathology and dysconnectivity in the vHIP-mPFC pathway, that is, prior to the appearance of Aβ plaques and tau tangles. Seven genes (*Gpm6b, Slc38a5, Ccr5, Kcnj10, Ddah1, Gad1, Slc17a8*) were significantly differentially expressed in 3– and 6-month-old 3xTg mice, revealing alterations to gene expression that may potentiate early-stage disruptions to hippocampal-prefrontal connectivity and subsequent executive dysfunction.

## Results

### 3xTg mice displayed decision-making deficits in the 4-Choice Gambling Task

To assess executive function in an AD context, male 3xTg and control mice were trained to criterion on the 4-Choice Gambling Task (4CGT, figure 1), a rodent analogue of the Iowa Gambling Task (IGT) used to assess decision-making in the clinic. Mice began staged training at 2-3 months of age and were fully trained to criterion by 6-7 months of age (n= 5 controls, n= 7 3xTgs). The training performance of both groups was similar, with no significant difference in both the overall number of sessions required to reach criterion and the number of sessions required to reach criterion for each training stage. Similarly, during testing, the number of trials completed per session by 3xTgs was similar to controls. Additionally, the percentages of omissions, premature responses, and number of perseverant responses to the choice location or rest of the grid were similar between control and 3xTg mice. Therefore, 3xTg mice were able to learn and perform the task at a similar rate and engagement level to control mice (supplementary figure 1).

**Figure 1:**
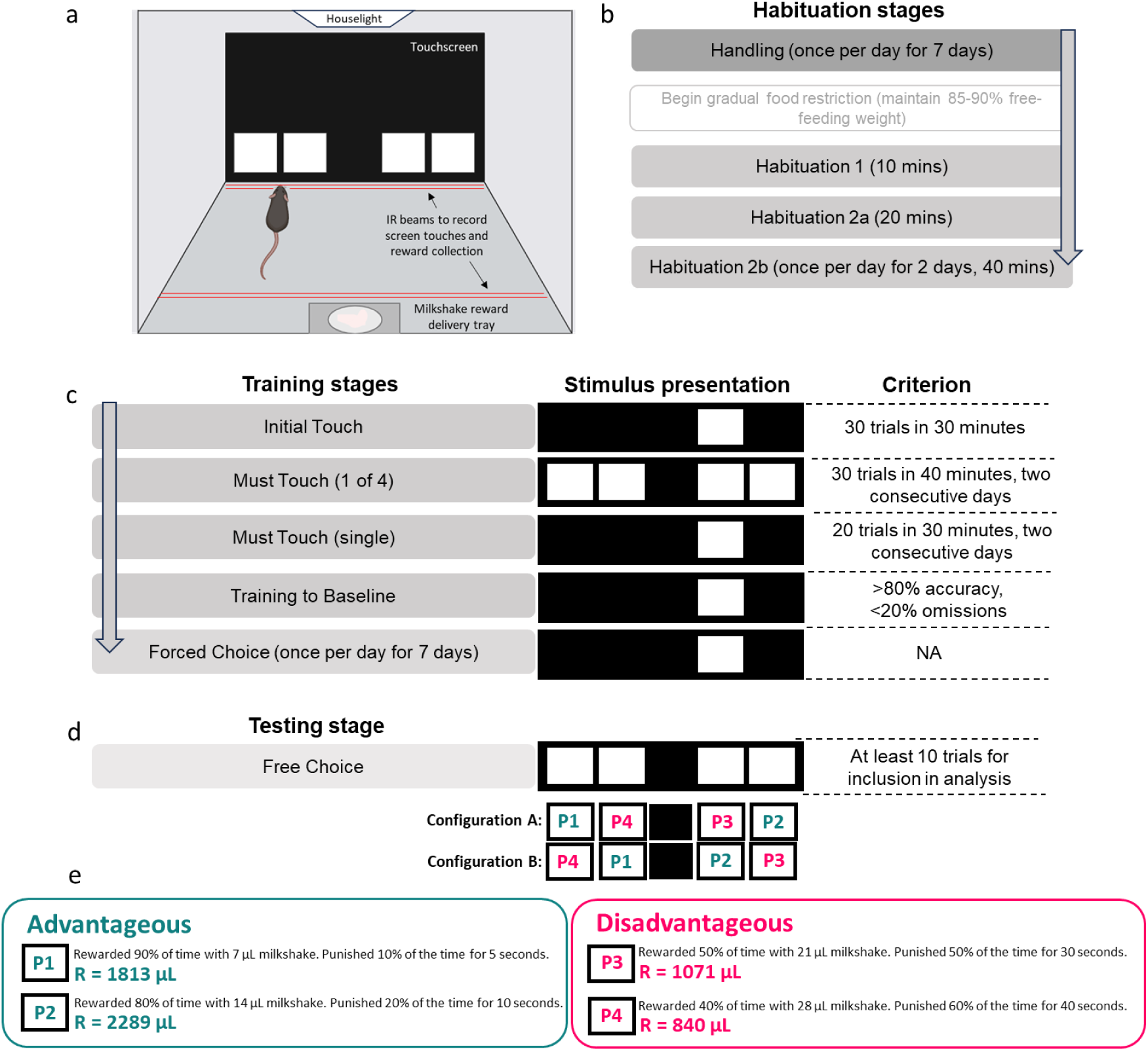
4-Choice Gambling task (4CGT) habituation, training, and testing procedures. **a** Set-up of the touchscreen-operant chamber in which the task is performed. **b** Order of 4CGT habituation stages. **c** Order of 4CGT training stages, their associated stimulus presentation and criterion required to pass each stage. **d** The 4CGT testing stage consisted of Free Choice sessions where mice were presented with all four choice stimuli. Two configurations were used, to prevent location bias. **e** Reward/punishment contingencies for each choice location. P1 and P2 were deemed low-risk, advantageous options, whilst P3 and P4 were high-risk, disadvantageous options. This can be deduced by the R-value, which represents the total volume of milkshake received over one 30-minute trial if that choice option is chosen exclusively throughout the trial.

Analysis of test sessions revealed no significant effect of session in control (figure 2a) or 3xTg (figure 2d) mice, nor a significant interaction between session and average choice %. However, there was a significant effect of average choice % in control mice (figure 2a, F (3,16) = 24.64; p<0.0001) but not 3xTg mice (figure 2d). When sessions were combined (4 sessions per block), the significant effect of average choice % remained (figure 2b, F (3, 16) = 29.37; p<0.0001). Control mice showed a clear preference for option P2, choosing it significantly more than other options in the first and second session blocks (figure 2b, p=0.0424 for P2 over P3 in first session block; p<0.0001 for P2 over P1 and P4, and p=0.0003 for P2 over P3 in second session block). Conversely, 3xTg mice did not choose P2 significantly more than other options even following session blocking (figure 2e). When combining advantageous versus disadvantageous options (P1+P2 versus P3+P4), control mice showed a clear preference for advantageous options (figure 2c, F (1,22) = 146.2; p<0.0001) and chose these significantly more than disadvantageous options in all three session blocks (p= 0.0010, p<0.0001, p= 0.0041) whilst 3xTgs did not show any preference (figure 2f).

**Figure 2:**
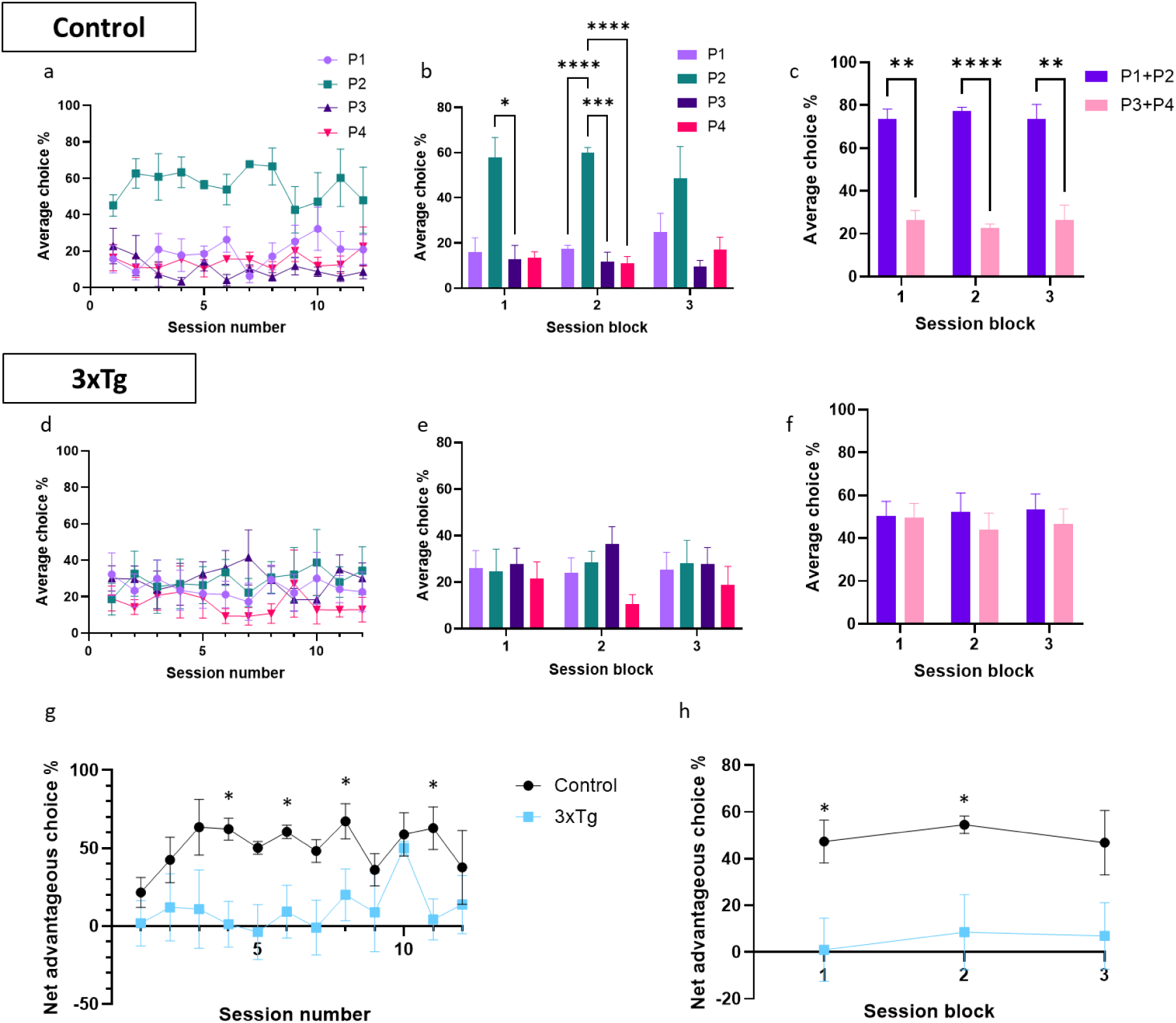
Performance of control and 3xTgAD mice on the 4-Choice Gambling Task. **a** Average choice % of control mice by session. **b** The same as for **a**, but with sessions displayed in blocks of 4. **c** Average choice % of control mice for advantageous (P1+P2) versus disadvantageous (P3+P4) options, by session block. **d** Average choice % of 3xTg mice by session. **e** The same as for **d**, but with sessions displayed in blocks of 4. **f** Average choice % of 3xTg mice for advantageous (P1+P2) versus disadvantageous (P3+P4) options, by session block. **g** Net advantageous choice % by individual session. This was calculated by subtracting the average % of disadvantageous choices (P3+P4) from the average % of advantageous choices (P1+P2) selected by mice across the session. The higher the value, the better the decision-making and task performance. Control mice had a significantly higher net advantageous choice % than 3xTg mice. **h** The same as for **g**, but with sessions blocked. Control mice had a significantly higher net advantageous choice % than 3xTg mice. n= 5 controls, n= 7 3xTgs. Error bars indicate SEM.

Performance on the 4CGT can also be assessed by comparing the net advantageous choice of control and 3xTg mice (24). This was calculated by subtracting the average % of disadvantageous choices (P3+P4) chosen over the course of the session from the % of advantageous choices (P1+P2) chosen. Mixed-effects analysis showed a significant effect of genotype on net advantageous choice; control mice had a significantly higher net advantageous choice across individual sessions, averaging 50.93% across 12 sessions, compared to 10.61% for 3xTg mice (figure 2g, F (1,10) = 8,203; p=0.0168). When sessions were blocked, this significant effect of genotype remained (figure 2h, F (1, 10) = 9.419; p=0.0119). Multiple comparison analysis showed that controls had a significantly higher net advantageous choice in individual session blocks 1 and 2 (figure 2h, p=0.0220 and p=0.0289, respectively). Control mice were, therefore, able to establish an advantageous strategy over the first few trials, and then maintain this strategy for subsequent sessions, whereas 3xTgs could not develop an advantageous strategy (figure 2g). As other aspects of task performance were similar between groups (supplementary figure 1), the inability of 6-8-month 3xTg mice to perform the 4CGT was most likely due to cognitive dysfunction and deficits in decision-making, as opposed to a lack of motivation, atypical locomotor activity, or an inability to learn the task.

### Altered input/output and paired pulse responses in the prefrontal cortex of 3xTg mice following stimulation of hippocampal inputs

To examine how vHIP-mPFC connectivity may be altered in 3xTg mice, local field responses were recorded in brain slices from layers II/III and V of the IL cortex following stimulation (10 to 100V intensities) of hippocampal fibres innervating these regions (52) (figure 3a). Representative responses for control and 3xTg mice are shown in figure 3b and 3d, respectively. In IL layer II/III, input-output curves showed a significant effect of genotype (p<0.0001) and stimulation strength (p<0.0001) and a significant interaction (figure 3c, F (9,150) = 3.658; p=0.0004). Pairwise comparisons showed that local field responses in IL layer II/III were significantly lower in 3xTg than control mice between 40 and 100V (figure 3c, p=0.0432, p=0.0053 and p=0.0007 for 40V, 50V and 60V stimulations respectively, p<0.0001 for 70V to 100V stimulations). Additionally, population spikes were more frequently observed in layer II/III of control animals compared to 3xTgs (figure 3f, F (3,129) = 92.06; p <0.0001). These observations support a reduced synaptic connectivity between the vHIP and mPFC, and dampened spread of postsynaptic excitation within the mPFC of 3xTgs. However, the presence and latency to maximal amplitude for a slower (presumed inhibitory) response component in layer II/III were similar between control and 3xTg LFPs (figures 3g and 3h). Response amplitudes in IL layer V were similar between 3xTgs and control mice (figure 3e).

**Figure 3:**
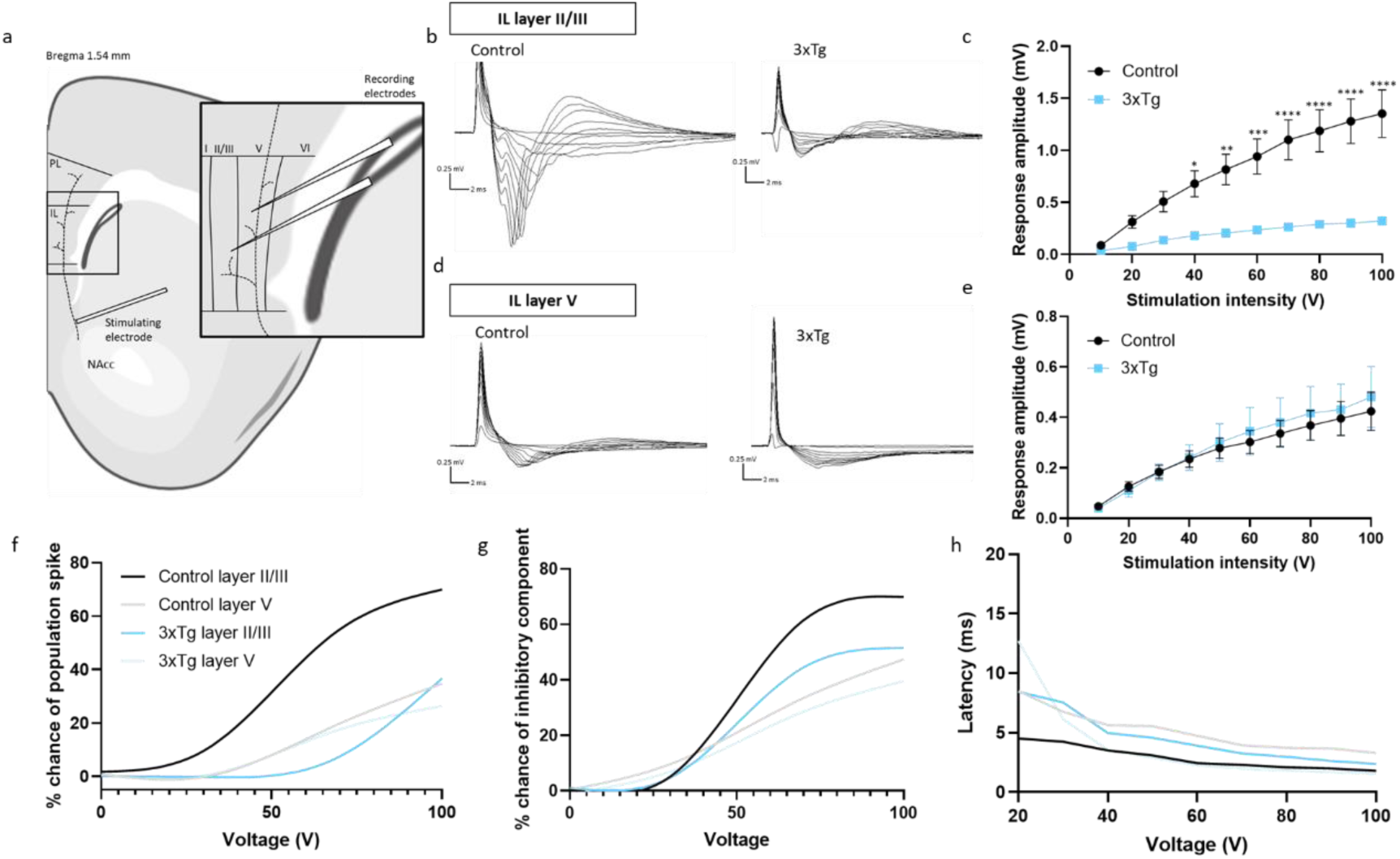
Input/output responses of 3xTgAD and control mice in mPFC layers II/III and V following hippocampal stimulation. **a** Schematic showing electrophysiology set-up. The stimulating electrode was placed alongside and dorsal to the nucleus accumbens core (NAcc), in the area reported to contain hippocampal fibres innervating the infralimbic (IL) cortex of the mPFC. Two recording electrodes were placed in the mPFC, in IL layers II/III and V, respectively, to record local field potentials. **b** Representative traces obtained from IL layer II/III of control (left) and 3xTg (right) slices at 10-100V stimulating strengths. **c** Input/output curve comparing response amplitudes in layer II/III of control versus 3xTg mice. **d** Representative traces obtained from IL layer V of control (left) and 3xTg (right) slices. **e** Input/output curve comparing response amplitudes in layer V of control versus 3xTg mice. **f** % chance of observing a population spike in control versus 3xTg mice in layers II/III and V. **g** % chance of observing a late inhibitory postsynaptic potential (IPSP) response in control versus 3xTg mice IL layers II/III and V. **h** Latency to maximum response amplitude in control versus 3xTg mice IL layers II/III and V. Data from one slice per animal, n= 9 controls and n= 8 3xTgs. Error bars indicate SEM.

Next, to further explore the synaptic short-term plasticity of hippocampal-prefrontal synapses, responses were recorded following paired pulse stimulation of innervating hippocampal fibres at 20, 50, 100, 200, and 500ms intervals. As with IO responses, paired pulse responses generally exhibited an initial excitatory local field response, followed by a slower presumed inhibitory component. In layer II/III, control mPFC LFPs exhibited mild paired pulse facilitation (PPF) at all intervals except 20ms, whereas 3xTg LFPs displayed a PPR very close to 1 (i.e., no facilitation) at all intervals except 20 ms, at which they exhibited paired pulse depression (PPD). However, 2-way ANOVA analyses of PPRs in layer II/III did not show a significant effect of genotype or interval, nor a significant interaction (figure 4a and 4b). Conversely, in layer V, a significant effect of genotype (F (1,50) = 40.79; p<0.0001) and a significant interaction between genotype and interval was observed (figures 4c and 4d, F (4,50) = 3.756; p=0.0095). Multiple comparisons analysis showed that PPRs at shorter intervals (20, 50 and 100ms) were significantly reduced in 3xTg compared to control mice (figure 4d, p<0.0001, p=0.0002 and p=0.0498 respectively). At all five intervals, 3xTg responses exhibited PPD, indicative of a reduction in short-term synaptic plasticity in the vHIP-mPFC layer V input in 3xTg mice.

**Figure 4:**
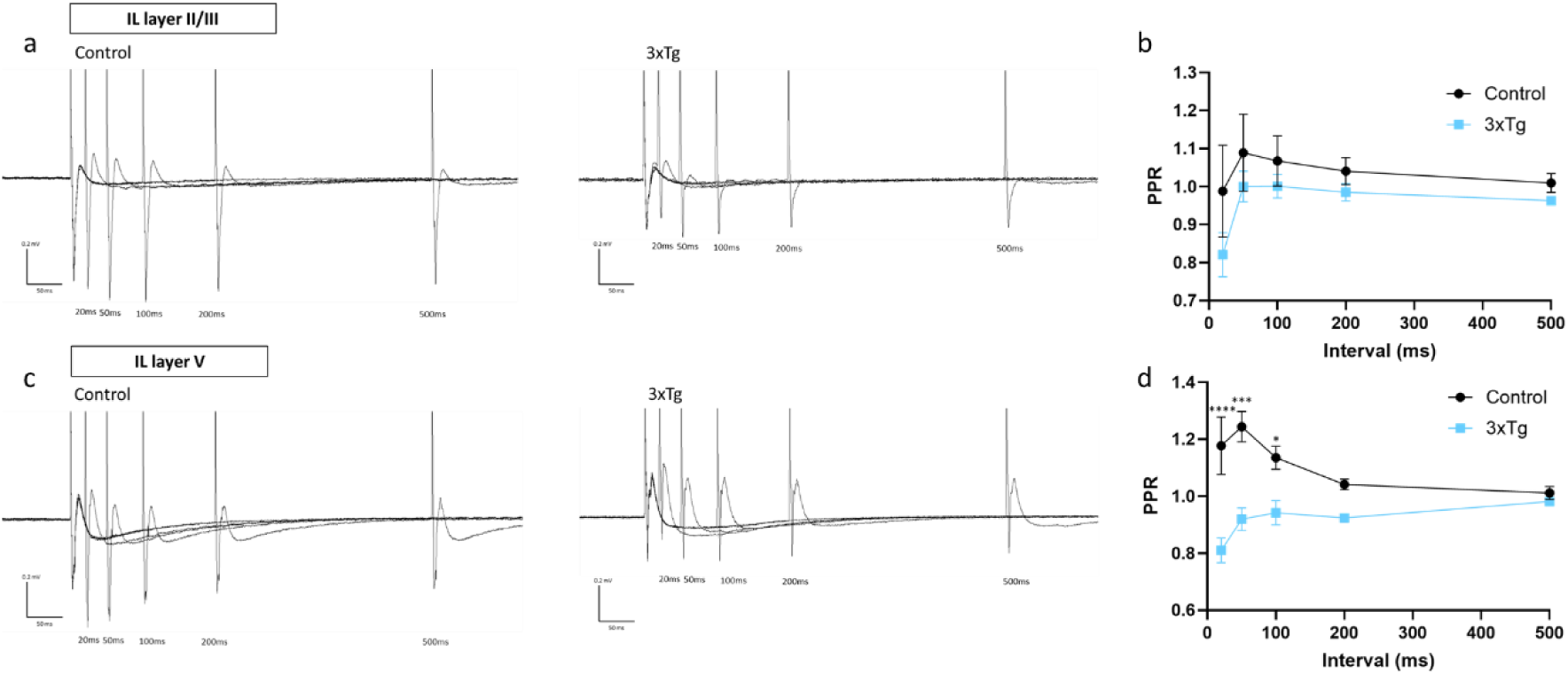
Responses of 3xTg and control mice in mPFC layers II/III and V following 100V hippocampal paired pulse stimulation at 20, 50, 100, 200 and 500ms. **a** Representative traces obtained from IL layer II/III of control (left) and 3xTg (right) slices. **b** Quantification of paired pulse ratios at each interval in layer II/III of 3xTgs and controls. **c** Representative traces obtained from IL layer V of control (left) and 3xTg (right) slices. **d** Quantification of paired pulse ratios at each interval in layer V of 3xTgs and controls. Data from one slice per animal, n= 9 controls and n= 5 3xTgs. Error bars indicate SEM

### 3xTgs displayed gene expression changes associated with glutamatergic, GABAergic, and dopaminergic activity, and calcium ion transport

To determine whether the behavioural and electrophysiological changes described above were associated with any significantly enriched genes at similar preclinical stages, bulk RNA sequencing analyses of the mPFC was performed in 3– and 6-month-old 3xTg and control male mice. Genes were grouped using the gene ontology (GO) major bioinformatics classification system, which contains cellular component, molecular function, and biological process branches. In general, gene expression changes were not as numerous or large in 3-month compared to 6-month animals, and no descriptive terms were significantly enriched in 3-month mice; figure 5a displays the number of genes associated with each term (the ‘Count’), the ratio of the number of differential genes linked with the term to the total number of differential genes (the ‘Gene ratio’), and the significance between the two groups given by the adjusted p-value (padj). Individual gene expression changes associated with neuronal synaptic function and neurotransmitter release were of particular interest in this study, as these are likely to underlie alterations to IO response amplitudes and PPRs. Seven genes of interest associated with these terms were identified as differentially expressed genes (DEGs) between 3xTg and control mice at 3 months of age: *Gpm6b*, *Slc38a5*, *Ccr5*, *Kcnj10*, *Ddah1*, *Gad1* and *Slc17a8/VGLUT3*. When plotted in a clustered heatmap, high levels of hierarchical clustering were observed between 3xTg and control mice for these genes (figure 5b). Several of these are associated with glutamatergic activity and transport, including *Slc38a5* (SNAT5), *Slc17a8* (VGLUT3), *GAD1* and *Kcnj10*, whilst *Gpm6b* and *Ddah1* have been linked with neurotransmitter activity and metabolism. *Ccr5* is a chemokine receptor that has been reported to play a role in calcium ion transport and glutamatergic function. The fold change of each of these genes in 3xTgs compared to controls is shown in figure 5c. Interestingly, substantial differences in the expression of genes associated with cytosolic calcium transport were not observed in 3-month-old animals, even though associated genes showed a high level of differential expression by 6 months. Thus, mPFC gene changes underlying attenuated glutamatergic synaptic transmission occur earlier in disease presentation than changes to expression of genes associated with calcium ion transport.

**Figure 5:**
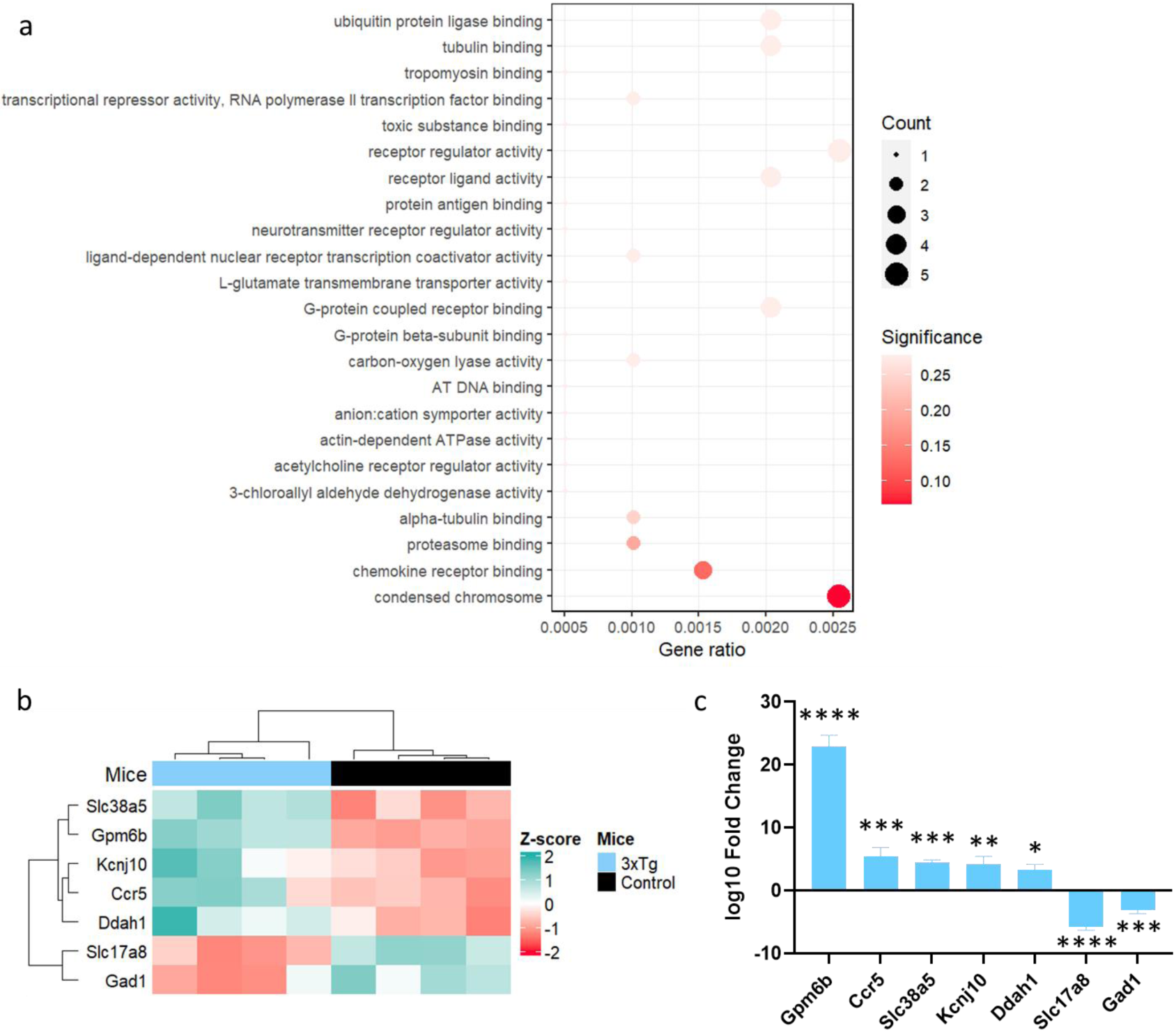
Early gene changes observed in the mPFC of 3-month male 3xTgAD and control mice. **a** Dot Plot showing most differentially expressed gene ontology (GO) terms between 3xTgs and control mice at 3 months and their respective adjusted p-value (padj), denoted by colour. The size of each dot represents the number of genes annotated to that specific term. The abscissa of each dot represents the ratio of the number of differential genes linked with that term to the total number of differential genes. No GO term was significantly enriched between 3xTgs and controls at 3 months. **b** Clustered heatmap showing individual genes included in terms relating to neurotransmitter and synaptic function which were significantly differentially expressed in both 3– and 6-month 3xTgs compared to controls. **c** Fold change of the differentially expressed genes shown in **b**, in 3xTgs compared to controls. n= 4 controls and n= 4 3xTgs. Error bars indicate SEM

More substantial gene expression changes were observed between 6-month-old control and 3xTg animals. Descriptive terms with most significant gene changes were related to immune function and defence responses, specifically the production of interleukins-1 and 6, tumour necrosis factor and MAPK and ERK1/2 kinase activity (figure 6a). Significant gene changes were also observed in several descriptive terms linked with AD, such as ‘response to lipoprotein particle’, ‘divalent metal ion transport’, and ‘negative regulation of neurogenesis’ (figure 6b). When terms associated with neuronal synaptic function and neurotransmitter release were compiled (figure 6c) and gene expression between controls and 3xTgs compared, terms associated with glutamatergic and GABAergic synaptic transmission, neurotransmitter function and transport, and calcium ion transport were significantly enriched (figure 6d). Genes associated with these descriptive terms were plotted on a clustered heatmap (figure 6e), which revealed high levels of hierarchical clustering between 3xTg and control samples, indicating large variability between control and 3xTg mice in the gene expression present within these descriptive terms.

**Figure 6:**
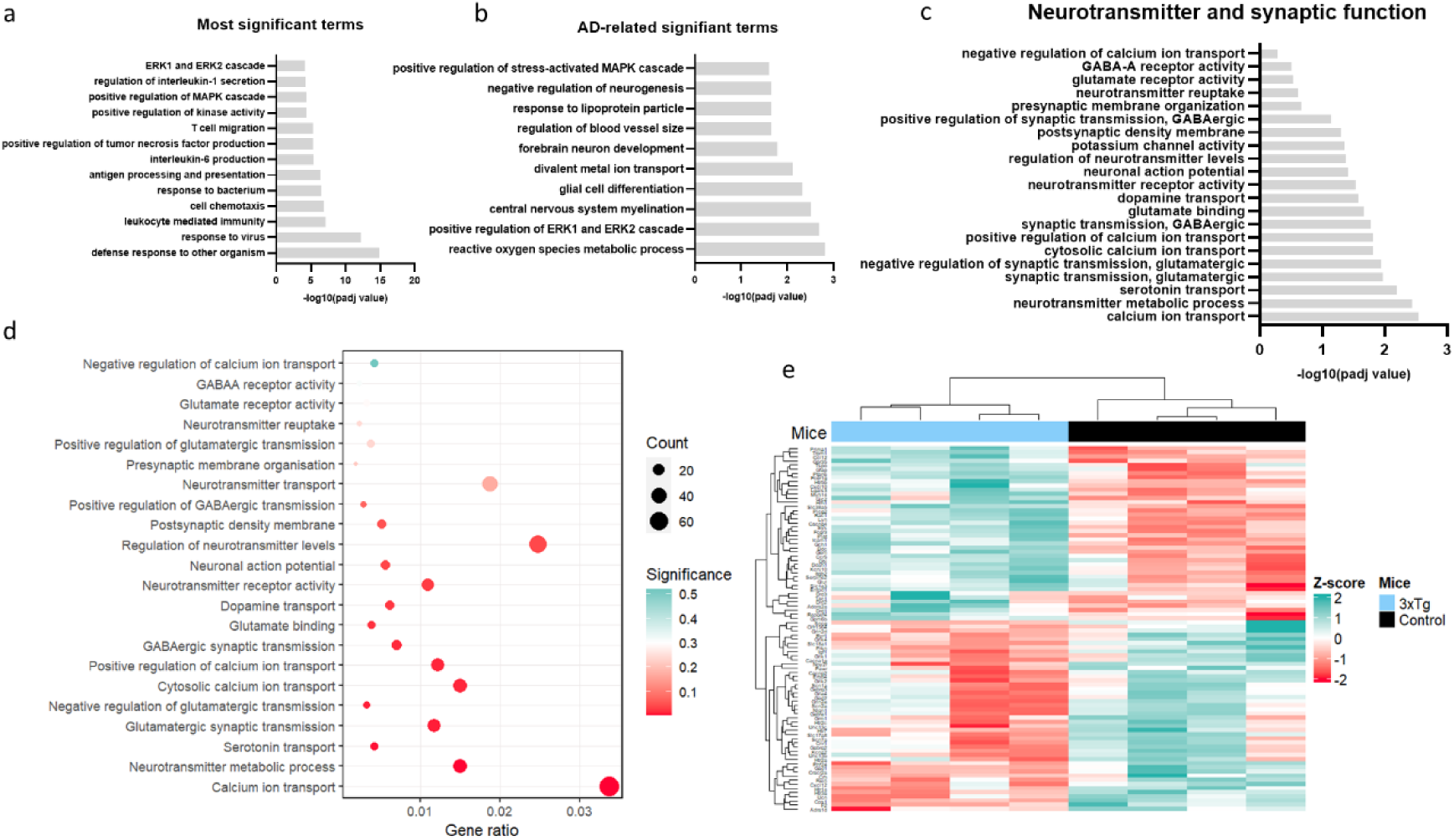
Significantly enriched GO terms identified by bulk mPFC RNA sequencing in 6-month-old male 3xTgs versus controls. **a** Most significant gene ontology (GO) terms of differentially expressed genes (DEGs) in the mPFC of 6-month-old 3xTgs compared to controls, and their adjusted p-value (padj). **b** Significant GO terms of DEGs relating to AD pathology and their padj. **c** GO terms of interest relating to neurotransmitter and synaptic function and their padj values. **d** Dot Plot showing the terms in **c** and their respective padj values, denoted by colour. The size of each dot represents the number of genes annotated to that specific term. The abscissa of each dot represents the ratio of the number of differential genes linked with that term to the total number of differential genes. **e** Clustered heatmap showing the expression pattern of genes included in the terms relating to neurotransmitter and synaptic function. n= 4 controls and n= 4 3xTgs.

Next, specific gene expression changes associated with the above significant terms in 6-month animals were investigated in more detail. Terms relating to glutamatergic function, including ‘glutamate binding’ (padj=0.0222), ‘glutamatergic synaptic transmission’ (padj=0.0175) and ‘negative regulation of glutamatergic synaptic transmission’ (padj=0.0113) were significantly enriched between groups. In 3xTgs, genes encoding for glutamatergic receptor subunits, such as *Grin2d*, *Grik1*, *Grin2a* and *Gria4* were downregulated, as was *Cps1* associated with glutamate binding (figure 7a). Genes associated with glutamatergic synaptic transmission, such as *Slc17a8* (encoding the vesicular glutamate transport 3; VGLUT3), *Prkn* and *Unc13c* were significantly downregulated, whilst *Adora2a*, *Plat*, *Drd1* and *Drd2* were upregulated (figure 7b). Genes associated with negative regulation of glutamatergic synaptic transmission, such as *Npy2r*, *Adra1d*, *Htr2a* and *Grik1/2,* were downregulated (figure 7c).

**Figure 7:**
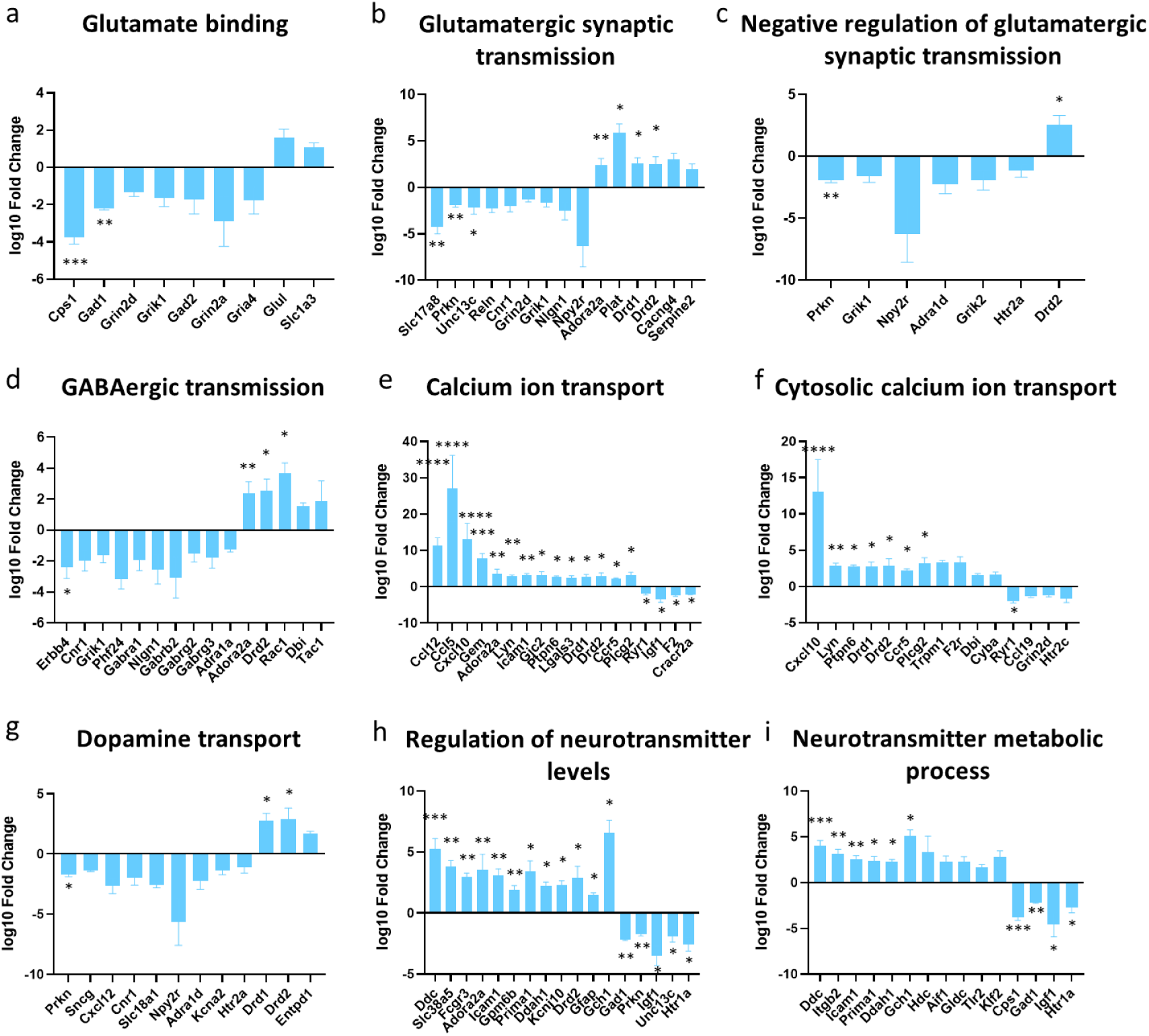
Changes to gene expression in 6-month-old male 3xTgs compared to controls, determined by bulk mPFC RNA sequencing. **a-c** Expression level in 3xTgs compared to controls of genes included in GO terms relating to glutamatergic function. **d** Gene fold change of genes relating to GABAergic function in 3xTgs compared to controls. **e, f** Gene changes relating to calcium ion transport in 3xTgs compared to controls. **g** Gene fold change of genes relating to dopaminergic function in 3xTgs compared to controls. **h, i** Gene changes relating to neurotransmitter function in 3xTgs compared to controls. n= 4 controls and n= 4 3xTgs. Error bars indicate SEM.

Changes to gene expression associated with GABAergic transmission were also plotted, to establish how changes to GABAergic function underlie alterations to the E-I balance. ‘GABAergic transmission’ was a significantly enriched term (padj=0.0165), and expression changes were observed in multiple genes including *Erbb4*, *Phf24*, *Nlgn1* (downregulated), *Rac1* (upregulated) and receptor subunits including *Gabra1*, *Gabrb2*, *Gabrg2*, *Gabrg3*, *Adra1a*, *Adora2a* and *Drd2*, most of which were downregulated in 3xTgs (figure 7d). Due to the frequent observation of gene expression changes to neurotransmitter receptor subtypes, further terms associated with neurotransmitter receptor function were also investigated, and ‘dopamine transport’ (padj=0.0265) ‘neurotransmitter receptor activity’ (padj=0.0296), and ‘neurotransmitter metabolic process’ (padj=0.00367) were all significantly enriched descriptive terms in 6-month-old 3xTg mice (figure 7g-i). Significant gene expression changes associated with these terms included the downregulation of *Prkn* and upregulation of *Drd1* and *Drd2* in 3xTgs.

Finally, to identify underlying gene changes associated with increased vesicle release probability in 3xTgs, as was observed in paired pulse stimulation experiments, gene expression changes underlying calcium ion transport were plotted. ‘Calcium ion transport’ (padj=0.00284) and, more specifically, ‘cytosolic calcium ion transport’ (padj=0.0152) were significantly enriched terms. Individual gene changes revealed a general, strong upregulation of genes involved in regulating calcium ion transport in both neurons and glia, including *Ccl5*, *Cxcl10*, *Lyn*, *Icam1*, *Gem* and *Ifg1* (figure 7e, f). Descriptive terms linked with vesicle release machinery, including ‘presynaptic membrane organisation’, ‘presynaptic membrane’, ‘synaptic vesicle fusion to presynaptic active zone membrane’ and ‘presynaptic process involved in chemical synaptic transmission’ were not significantly enriched. This suggests that observed alterations to the vesicle release probability in 3xTgs are due to changes in calcium ion transport in the presynaptic membrane, as opposed to variable expression of proteins associated with vesicle release.

## Discussion

Executive dysfunction is a marked feature of dementia and AD; however, our understanding of this symptom realm is lacking, due to relatively little research focus in preclinical models of dementia. Alongside control mice, 6-month-old male 3xTgAD mice were able to successfully train to criterion on the 4-Choice Gambling Task. Subsequent testing revealed that, whilst control mice were able to optimise their strategy and choose advantageous (low risk) options a significantly higher proportion of the time than disadvantageous (high risk) options, 3xTg mice did not show preference for advantageous options. Indeed, 3xTg mice displayed similar preference for all reward options (increased disadvantageous choices compared to control), suggesting an inability to adapt behaviour based on outcomes (reward prediction error). This profile closely resembles the performance of patients with mild AD on the IGT, the clinical equivalent of the 4CGT. (24). Such poor decision-making on the IGT may arise due to either substantial risk-taking behaviour whilst the task is being undertaken ‘under risk’ (subjects continuously choose high-risk, high reward options) or the inability to develop an advantageous strategy whilst the task is being played ‘under ambiguity’ (subjects switch frequently between known advantageous and disadvantageous options without an obvious strategy) (24). Similarly to IGT results in AD patients, 3xTg mice did not choose predominantly high-risk, high-reward choices, and instead chose a similar percentage of advantageous and disadvantageous options, thereby revealing that poor performance on the task was most likely due to the inability to plan and implement an appropriate strategy. This form of executive dysfunction in AD patients has been attributed to altered function of the dorsolateral prefrontal cortex (21,24,53,54), analogous to the rodent mPFC. Here, hippocampal input to the mPFC would normally play a vital role in supporting memory encoding by continuously updating task-relevant information to inform decision-making (55). The hippocampal-prefrontal pathway is, therefore, essential in enabling prefrontal neurons to compute which actions are appropriate in a given spatial context, using memories of that context encoded by the hippocampus (56). The strengthening of this pathway, observed by theta-frequency connectivity, has been strongly suggested to enable learning (2), whilst lesions of the vHIP abolish anticipatory activity in the mPFC, altering impulse control and goal-directed behaviour (57). The latter results in the inability to utilize task-relevant information for the formation of an appropriate strategy. This choice profile contrasts to examples of risk-taking behaviour where subjects cannot stop choosing high risk, high reward options, as observed in patients with amygdala or ventromedial PFC lesions (58,59). The present behavioural results therefore implicate deficits in vHIP-mPFC connectivity and function in the loss of executive function commonly observed in AD patients (3,12–15).

### Input-output response amplitudes

To determine whether the behavioural outcomes described above were indeed associated with altered hippocampal input to the mPFC, we examined the physiology of this synaptic connection in our 3xTg AD model *in vitro*. Pyramidal neurons in the mPFC have previously been shown to display reduced neuronal activity in other AD mouse models (60,61) and the same phenotype was observed in this study, specifically in 3xTg prefrontal neurons receiving direct hippocampal input. Prefrontal local field responses to hippocampal fibre stimulation have been reported to display an initial excitatory phase, reflecting positive ion influx (known as the excitatory postsynaptic current, EPSC) that leads to fast AMPA-receptor and slow NMDA-receptor activation in the form of an excitatory postsynaptic potential (EPSP) (9,62,63). This is sometimes succeeded by a population spike. The initial excitatory response is followed by an inhibitory postsynaptic potential (IPSP), as a result of GABA_A_ and GABA_B_ receptor activation, which leads to chloride ion influx and potassium ion efflux (underlying the inhibitory postsynaptic current, IPSC) (9,64). Therefore, input-output response amplitudes depend on the level of activation of both glutamatergic and GABAergic receptors.

In this study, following single pulse stimulation of hippocampal input fibres in 3xTgs, a significant reduction in mPFC local field response amplitude was observed compared to controls in mPFC layer II/III, but not layer V. Additionally, mPFC LFPs were less likely to display a population spike in layer II/III of 3xTg mice. Therefore, by recording LFPs in the mPFC following hippocampal fibre stimulation, the connectivity between the hippocampus and layer II/III mPFC of 3xTg mice was shown to be compromised, leading to a reduced E-I balance within the mPFC due to a general increase in inhibition, and/or a reduction in excitation. Local mPFC circuitry involving interneurons has been shown to be layer specific; layer II/III interneurons receiving hippocampal input predominantly express vasoactive intestinal polypeptide (VIP), whilst those in layer V express parvalbumin (PV), somatostatin (SOM) or cholecystokinin (CCK) (65–68). Layer V interneuron subtypes have been suggested to contribute towards feedforward and feedback inhibition by synapsing directly with excitatory pyramidal cells in layer V (65,69,70). Conversely, VIP-expressing interneurons inhibit other interneuron subtypes in layer II/III (8), thus, leading to disinhibition. Observed layer-specific differences in the response amplitude may, therefore, be associated with differences in local interneuron circuitry. Previous studies have reported changes to firing properties of VIP-positive interneurons in the hippocampus in young 3xTg mice (71), and during ageing (72), although whether alterations to VIP-positive interneurons also occur in layer II/III of the mPFC, and how this may underlie alterations to the E-I balance, have yet to be determined.

#### Multiple gene changes may contribute to reduction in IO response amplitudes in 6-month-old 3xTgs

Reductions in the excitatory/inhibitory balance may occur as a direct result of increased GABAergic inhibitory transmission, or attenuated glutamate receptor expression and synaptic transmission, and thus reduced glutamatergic binding at the postsynaptic membrane. RNA sequencing data highlighted several gene changes associated with glutamate binding and transport and GABAergic signalling in 6-month-old animals. Gene changes linked with glutamatergic function in 3xTgs include the downregulation of *Cps1*, *Gad1*, *Slc17a8* and *Unc13c*.

Although not significant, DEGs *Grin2d*, *Grik1*, *Grin2a* and *Gria4*, which encode glutamate receptor subunits, were all downregulated in 6-month-old 3xTgs compared to controls and contributed towards significant enrichment of ‘glutamate binding’. Additionally, 6-month-old 3xTg mice displayed gene changes associated with GABAergic transmission, including *Erbb4* (73,73,74), *Adora2a* (75,76) and *Rac1* (77,78), which may also underlie disruptions to the maintenance of the E-I balance.

Given its reported role in the modulation of glutamatergic activity associated with hippocampal-prefrontal plasticity (79,80), gene changes associated with dopamine transport in both 3– and 6-month-old animals were noteworthy. By 6 months, ‘dopamine transport’ was a significantly enriched term, and gene expression changes in 3xTgs relating to this term were observed. These included downregulation of the Parkin gene (*Prkn*), and upregulation of dopamine receptors 1 and 2 (*Drd1,2*). Increased expression of *Prkn*, which is strongly associated with familial Parkinson’s disease, has been reported to increase dopamine uptake by upregulating expression of the dopamine transporter (81). Expression of dopamine receptors 1 and 2 (D1 and D2 receptors) in the PFC are considered critical for behavioural flexibility (82), reward-driven behaviours (83), and learning and memory processes dependent on synaptic plasticity (2,84,85). D1 receptors have been shown to colocalise with glutamate receptors in PFC pyramidal cells (86), and both D1 and D2 receptors directly modulate glutamate receptor function and transmission, and thus synaptic plasticity (87–90). Interestingly, the application of a D2 receptor antagonist has been shown to improve performance on the 4CGT in male rats (25), suggesting that overactivation of D2 receptors may contribute towards impaired decision-making.

#### Candidates for early drivers of disease pathology identified by RNA sequencing in 3-month-old 3xTgs

*Genes involved in the maintenance of the E-I balance.* The *Kcnj10* gene encoding for the inward-rectifying potassium channel 4.1 (Kir4.1) has been linked with alterations to neuronal excitability; activation of the channels on astrocytes acts to elevate extracellular potassium and glutamate (91). Furthermore, Kir4.1 has previously been linked with aberrant neuronal activity in AD, specifically in the prefrontal cortex of 3xTg mice (92). In the present study, *Kcnj10* was found to be upregulated in both 3– and 6-month-old 3xTgs, further implicating its altered expression in AD pathogenesis, in this case in mPFC neurons receiving hippocampal input.

*Slc38a5* encodes for the microglial solute neutral amino acid transporter 5 (SNAT5). This transporter has been suggested to regulate the glutamate-glutamine cycle (93), and the expression of NMDA receptors dependent on extracellular levels of glycine (94), thereby impacting glutamatergic transmission. This gene was upregulated in 3-month-as well as 6-month-old 3xTgs.

Of the gene changes associated with glutamatergic function, *Gad1* and *Slc17a8* were also significantly differentially expressed in 3-month 3xTgs. *Slc17a8* encodes the vesicular glutamate transport 3 (VGLUT3), an essential component of glutamate transport into synaptic vesicles. Reduced expression of VGLUT1 and 2 has been observed in prefrontal regions of AD patients, correlating with cognitive decline (36). The role of VGLUT3 has been relatively understudied in an AD context, although one recent report has suggested that loss of VGLUT3 negatively impacts learning and memory in mice (95). *Gad1* encodes glutamate decarboxylase 1 (GAD1), which catalyses the production of GABA from glutamate. The downregulation of PFC GAD1 expression inevitably disrupts the glutamate-glycine-GABA cycle and has been suggested to contribute towards E-I network and working memory impairments in schizophrenia (96). Reduced *Gad1* expression has previously been observed in AD patients (35,97).

*Genes involved in the neuromodulation of glutamate and GABA.* At 3 months, 3xTgs displayed significant upregulation of the *Ddah1* gene encoding for dimethylarginine dimethylaminohydrolase 1 (DDAH1). DDAH1 is a regulator of nitric oxide synthase and therefore plays a role in nitric oxide production (98). Overproduction of nitric oxide has been widely reported to contribute to neurotoxicity associated with AD due to its ability to exert damage in free radical form (99). Overexpression of DDAH1 has been shown to attenuate oxidative damage and reduce the secretion of Aβ in human cell cultures expressing APP_swe_ (100). Therefore, its upregulation in 3xTg mice may be a compensatory mechanism. The enzyme has also been reported to influence dopamine metabolism; its knock-out decreased dopamine metabolite concentration in the piriform cortex and striatum, impacting mouse exploratory behaviour (101). It is unknown how altered expression of this enzyme may affect dopamine production in the PFC, and how this influences excitability in the hippocampal-prefrontal pathway.

A role for the *Gpm6b* gene, encoding for the neuronal membrane glycoprotein M6-b which is widely expressed in both neurons and glia (102), in neuronal function is yet to be fully determined, although it has been suggested to alter signalling associated with serotonergic transmission (103), proposing another possible mechanism for aberrant neuromodulatory activity. A role for altered serotonergic activity in AD has been proposed in several recent reports (104–106).

### Paired pulse ratios

Short-term plasticity changes in the vHIP-mPFC pathway were explored via paired pulse recordings. PPF in prelimbic and infralimbic regions following hippocampal paired-pulse stimulation at intervals between 50 and 150ms have previously been observed, with largest PPF reportedly occurring at an interval of 50-75ms (107). A PPR less than 1 indicates that the first pulse is relatively larger than the second, and this is caused by increased vesicle release probability and subsequent vesicle depletion associated with PPD and synaptic weakening. A lower vesicle release probability results in a PPR larger than 1, indicative of PPF and synaptic strengthening (108). Presynaptic vesicle exocytosis is triggered by the influx of calcium, which initiates the activation of vesicle release machinery, namely synaptotagmins, complexins and SNARE proteins (44). Therefore, changes to release probability can be explained by variations in the intracellular accumulation of residual calcium or functioning of vesicle release machinery.

In this study, significant reductions in the PPR were observed in mPFC LFPs of 3xTg mice compared to controls in layer V, but not layer II/III. It is possible that the lack of change in layer II/III is a consequence of the large reduction in response amplitudes following stimulation of excitatory hippocampal input in 3xTgs, which becomes insufficient to generate the levels of activation and recruitment of inhibition required to reveal changes to the PPR. In layer V, PPRs were significantly lower in 3xTgs, and exhibited PPD as opposed to the PPF observed in controls animals. These changes propose alterations to the vesicle release probability and/or residual calcium concentration of vHIP-mPFC synapses. These effects were particularly notable at shorter intervals (25, 50, and 100 ms), possibly indicating upregulation of inhibition mediated by GABA_A_ receptors.

#### Minimal changes to genes associated with calcium ion transport during early-stage AD

In 3-month-old 3xTgs, expression changes for genes underlying GABA_A_ receptor activity, synaptic vesicle release or calcium ion transport were only slight, aside from significant upregulation of the *Ccr5* gene which encodes for the C-C chemokine receptor 5 (CCR5). CCR5 activity has also been proposed as a modulator of glutamatergic transmission in glia-neuronal crosstalk (109), and CCR5-induced glutamate exocytosis has been observed in the spinal cord (110).

#### Extensive alterations to gene expression associated with calcium ion transport in 6-month-old 3xTgs

In 6-month-old animals, although significant changes associated with vesicle release machinery genes were not observed, genes linked to ‘cytosolic calcium ion transport’ were significantly differentially expressed, proposing alterations to calcium dynamics in 3xTgs. Expression of the *Cxcl10* gene encoding for chemokine ligand 10, which has been reported to increase neuronal cytosolic calcium release (111), was upregulated. Genes underlying calcium release into glia were also upregulated, including *Ccl5* (109) and *Lyn* (112), as was *Icam-1,* which has been reported to increase intracellular calcium in brain endothelial cells (113). Taken together, these gene changes propose drastic alterations to calcium dynamics in the brain parenchyma of 6-month-old 3xTg males compared to controls.

Due to the substantial number of gene changes associated with calcium ion dynamics, it is more likely that alterations to intracellular calcium dynamics, rather than vesicle release machinery, underlie modified vesicle release probability and, hence, PPD of 3xTg hippocampal-prefrontal synapses observed in paired-pulse recordings. Dysregulation of calcium ion dynamics has been linked with the accumulation of Aβ and is widely suggested to contribute towards aberrant synaptic signalling underlying neurodegeneration (47–50). An increased concentration of cytosolic calcium has also been linked to upregulated intraneuronal amyloid production, leading to cell death (114).

These results suggest that adjustments to the mPFC E-I imbalance and neuromodulation of glutamate and GABA occur in early stages of the disease. In contrast, disruptions to calcium ion transport and vesicle release probability emerge later in disease presentation, but still precede the appearance of Aβ plaques and tau tangles. It is possible that early changes to glutamatergic activity in layer II/III drive postsynaptic reductions in effective connectivity in the hippocampal-prefrontal pathway. This is counteracted by altered calcium transport at the presynaptic membrane, leading to increased vesicle release and thus PPD of hippocampal-prefrontal synapses in layer V. Additionally, alterations to gene expression associated with GABAergic transmission are observed by 6-months and may be another form of compensation for aberrant glutamatergic transmission. Changes to dopaminergic modulation of this pathway, specifically abnormal activity of D1 and D2 receptors, are likely to contribute towards these observations (1,90,115,116). To confirm an exact mechanism underlying observed changes, layer-specific single cell analyses (for example, using single cell RNA sequencing) will be required.

These results show that 3xTgAD mice can be trained to criterion on the touchscreen operant 4-Choice Gambling Task, and that this rodent version of the clinically utilized Iowa Gambling Task can be used successfully to observe executive dysfunction in an AD context. 3xTg mice displayed poorer performance on the task than matched controls, indicative of deficits in decision-making. Alongside these findings, altered mPFC responses to activation of hippocampal input fibres in 3xTg mice correlated with the identification of several genes displaying differential expression in the mPFC of 3xTgs which may contribute to their altered E-I balance and executive dysfunction, both symptomatic of early Alzheimer’s disease. Future studies may shed light on the precise neuronal type and the intricate neural networks, including dopaminergic, glutaminergic, and GABAergic pathways connecting to the mPFC region, that may be impaired in 3xTg mice.

**Supplementary figure 1:**
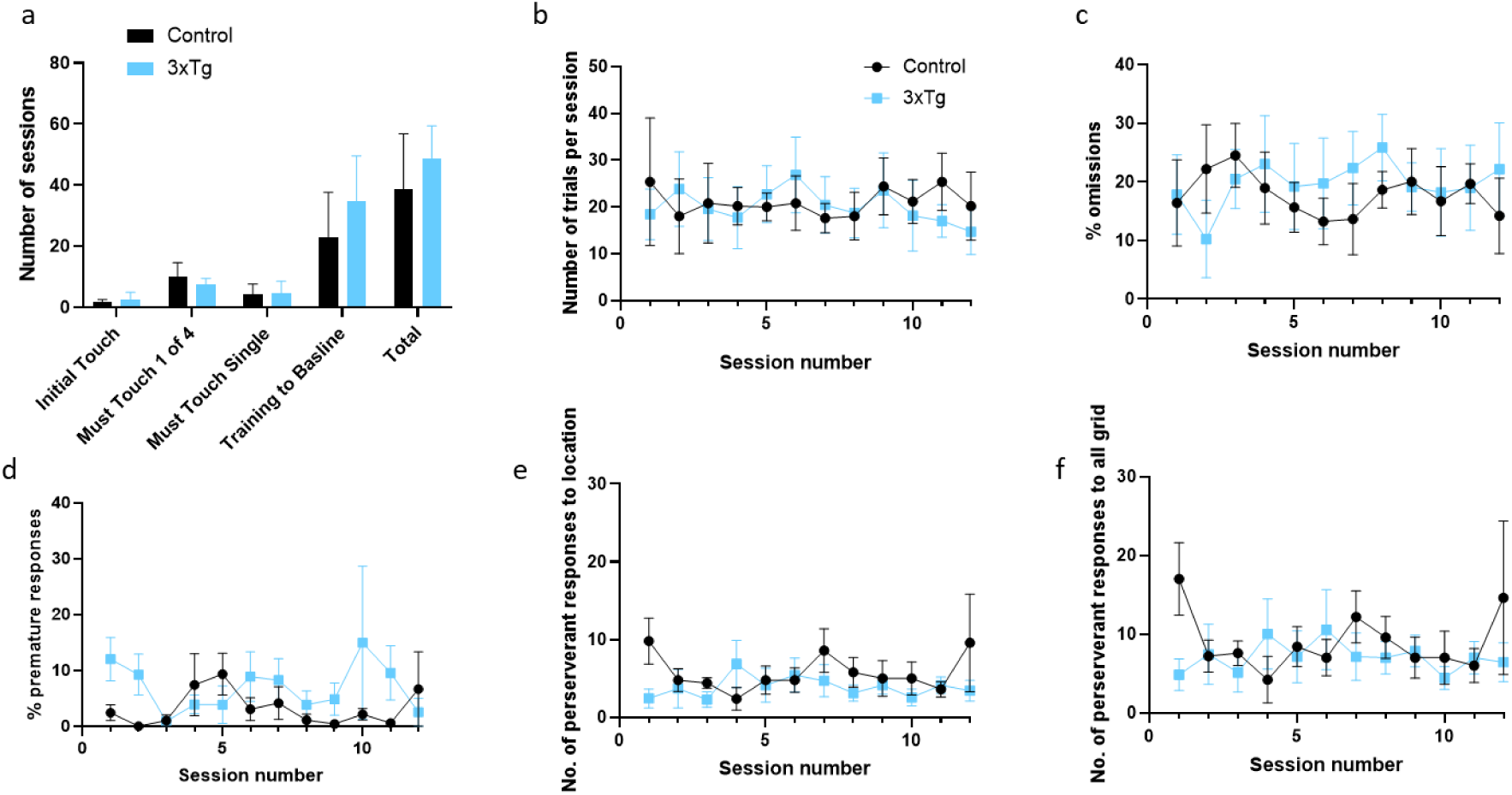
No significant differences in the sessions required to reach criterion, % omissions, % premature responses, or number of perseverant responses between 3xTg and control mice on the 4CGT. **a** Comparison between controls and 3xTgs in the number of sessions required to reach criterion at each training stage, and overall. **b** Comparison between controls and 3xTgs in the number of trials completed per session. **c-f** Comparisons between controls and 3xTgs in the **c** % omissions, **d** % premature responses, and number of perseverant responses to either **e** choice location or **f** all of grid, for each session. n= 5 controls and n= 7 3xTgs. Error bars indicate SEM.

## Materials and Methods

### Animals

*Behaviour and electrophysiology*. Male triple transgenic (3xTgAD; n= 9) mice carrying APP_SWE_, PS1_M146V_ and Tau_P301L_ transgenes, and matched controls (n= 9) of the same background strain (C57/129sv) were 2-3 months old at the start of behavioural training. At the start of behavioural testing, n= 7 3xTg and n= 5 control mice were 6-7 months old and had met pre-testing criteria. Mice that had not reached pre-testing criteria were removed from behavioural testing. For electrophysiology experiments, 3xTg (n= 8 and n= 5 for IO and PPR experiments, respectively) and control (n= 9) mice were 6 months of age, an approximate age-match to behaviourally tested mice. Mice were bred from homozygous pairs (Jackson Labs) in the University of Manchester Biological Services Facility. Genotyping of offspring mice was performed by Transnetyx using PCR amplification of tissue samples collected from ear punches. Mice were housed five per cage and maintained at a 12-hour light/dark cycle (lights off at 7pm). Mice had access to food and water ad libitum, except during behavioural training and testing, when they were food restricted to encourage behavioural performance whilst maintaining body weight no lower than 85-90% of free feeding animals. All experimental procedures were performed under a Home Office UK project licence in accordance with the Animals (Scientific Procedures) Act 1986, and approval by the University of Manchester Animal Welfare and Ethical Review Body (AWERB).

*RNA sequencing*. 3– and 6-month-old male triple transgenic (3xTgAD; n= 4 for each age) mice carrying APP_SWE_, PS1_M146V_ and Tau_P301L_ transgenes, and matched controls (n= 4 for each age) of the same background strain (C57/129sv) were used for bulk RNA sequencing of the medial prefrontal cortex. Mice were purchased from Jackson Labs and were housed under a 12-hour light/dark cycle in the Singapore A*STAR Biological Resource Centre with access to food and water ad libitum. All experimental procedures were approved by the Institutional Animal Care and Use Committee of the A*STAR of Singapore.

### Behavioural testing

*Behavioural apparatus.* Touchscreen-operant chambers (Campden Instruments Ltd., UK), as previously described (117–119), consisted of a trapezoidal floor surrounded by black Perspex walls, under which there was a waste-tray. The longest edge of the chamber contained the touchscreen, in front of which was a black Perspex mask containing five response windows. These windows corresponded to squares which would light up on the screen, with all but the middle square being used during 4CGT training and testing. The opposite side of the chamber contained a food delivery magazine and a light. Infra-red beams to measure task performance were installed in front of the touchscreen and the food reward trough (figure 1a). Eight chambers were used simultaneously within individual light– and sound-attenuating lockable boxes. These boxes also contained a house light, tone generator, ventilating fan, and camera. A peristaltic pump was used to deliver behavioural reinforcer (Yazoo strawberry milkshake, FrieslandCampina, Ltd., UK) into the food delivery magazine. ABET II software (Campden Instruments Ltd, UK) was used to control each touchscreen chamber. Training and testing protocols were performed according to the Mouse Touch 4C-GT ABET II Manual V1.2 (Campden Instruments Ltd., UK).

*4CGT habituation.* Prior to behavioural training, mice were habituated to experimenter handling every day for one week. They were then gradually food restricted for four days prior to the first training day to increase motivation for milkshake reward. Food restriction was continued throughout training and testing stages, to maintain motivation. Each mouse was first placed inside an experimental chamber for 10 minutes to allow habituation to the apparatus. On the second day of habituation, mice were left for 20 minutes, and milkshake reward (7 µl) was dispensed each time the mouse poked its head into the milkshake delivery tray. This was repeated on the third and fourth habituation days, which were 40 minutes long. By the fourth habituation day, all mice were able to successfully find and collect milkshake reward. Habituation stages are summarised in figure 1b.

*4CGT training.* Behavioural training (figure 1c) began immediately following habituation stages. Mice were given one daily training session, five days a week. The first stage was Initial Touch training, consisting of one white square being pseudo-randomly displayed in one of the four possible locations for 30 seconds. The square was then removed and 7 µl milkshake delivered (accompanied by a tone). If the square was nose poked during its display 20 µl milkshake was rewarded, encouraging touches to the stimuli. A new trial would be initiated once the mouse consumed the milkshake reward. Criterion at this stage was the completion of 30 trials in 30 minutes. The next stage required mice to touch the screen to receive milkshake reward; all four possible square locations were lit up and mice were rewarded with 7 µl milkshake for touching any 1 of the 4 squares. Criterion for this stage was 40 successful touches to the light cues within 30 minutes, on two consecutive days. The next stage was the same, except a single square only was pseudo-randomly presented across the four possible positions on a given trial, and milkshake dispensed only following touches to that target. Following the successful completion of 20 trials in 30 minutes on two consecutive days, mice were moved onto the final training stage. This was the same as the previous stage, except a time limit was imposed on the stimulus presentation, and mice were punished (luminance inverted for 5 seconds) for touches to incorrect/blank locations. This Training to Baseline stage contained up to 4 sessions, with the stimulus duration for each session lower than the one before (37, 21, 13 and 10 seconds respectively). Moving to the next session required mice to finish the previous session with >80% accuracy of touches to the correct stimuli and <20% trial omissions. Inclusion of this training stage is not deemed necessary (32), but the first Training to Baseline training stage (37-second stimulus duration) was included in this study to verify that mice could execute the task. For each training and testing stage, the number of trials completed, % accuracy and % omissions were recorded.

Finally, following training and prior to the Free Choice version of the task, mice were put through ten Forced Choice sessions where just one stimulus position was illuminated at once, to familiarise them completely with the reward/punishment possibilities for each position. Once they had successfully moved through all Forced Choice sessions, animals were tested on the Free Choice task. For both Forced Choice and Free Choice sessions, two spatial configurations of 4 positions were used to counterbalance possible location biases. Therefore, half the mice faced an order of choices of (from left to right); P1, P4, P2, P3 (configuration A), and the other half P4, P1, P3, P2 (configuration B, figure 1d).

*4CGT testing.* The Free Choice task began with 7 µl milkshake delivery, accompanied by a tone. Once the mouse nose poked the delivery magazine, it was presented with white square stimuli in positions 1, 2, 4 and 5. Mice had to touch one square, at which point they would receive either a milkshake reward or a time-out (the image would flash at a frequency of 5 Hz and chamber lights were inverted). Each square was associated with a different volume of milkshake reward, a different probability of receiving this reward instead of a time-out, and a different length of time-out if this occurred instead. Touches to squares with lower volumes of reward were also associated with a greater chance of reward, and shorter time-out length if the reward was not received. For example, touches to position 2 (P2) dispensed 14 μl of milkshake 80% of the time, whilst 20% of touches resulted in a 10 second time-out. Alternatively, touches to position 4 (P4) dispensed 28 μl of milkshake 40% of the time, and time-outs occurred for 60% of touches and were a much longer 40 seconds. A summary of all 4 positions’ reward/punishment contingencies can be seen in figure 1e. The order of best to worst choice in terms of highest long-term reward pay-off was calculated by comparing the maximum milkshake reward for each choice if that choice was chosen exclusively throughout the 30-minute task. This is referred to as the R value, with a higher value correlating with a larger volume of reward for that choice. The order of choices with highest to lowest R number was: P2 > P1 > P3 > P4. Therefore, the optimal volume of reward per unit time is achieved when the square at P2 is continuously chosen, and high-risk, high-reward options (P3 and P4) are avoided.

### Electrophysiology

*Brain slice preparation.* Mice were euthanised by cervical dislocation and brains quickly removed into ice-cold sucrose Krebs solution, which contained (in mM): 124 NaCl, 26 NaHCO3, 2 KCl, 1.25 KH2PO4, 10 MgSO4, 0.5 CaCl2, 10 D-glucose and 202 sucrose (290-295mOsm) and was continuously bubbled throughout slice preparation with 95% O_2_/5% CO_2_. Coronal slices (400 µM) were cut using the Vibroslice HA752 vibratome (Campden Instruments Ltd) in the same ice-cold solution, before they were transferred into holding chambers at 37°C containing artificial cerebrospinal fluid (aCSF). The aCSF solution consisted of (in mM): 124 NaCl, 26 NaHCO3, 2 KCl, 1.25 KH2PO4, 1.5 MgSO4, 1.5 CaCl2 and 10 D-glucose (290-295mOsm) and was continuously bubbled with 95% O_2_/5% CO_2_ throughout slice recovery and recordings. Slices were left to recover for at least 1 hour.

*Electrophysiological set-up.* Following recovery, slices were placed into a recording chamber, which was constantly perfused with 32°C aCSF at a flow rate of 0.7 ml/min. The recording chamber beneath the brain slice was filled with water, which was also bubbled with 95% O_2_/5% CO_2_ and heated to 32°C, ensuring a humidified, oxygen-rich atmosphere surrounded the slice. Once the slice had been placed in the recording area, a stimulating electrode was placed in the target area. Because the aim was to stimulate hippocampal fibres innervating the mPFC, the stimulating electrode was placed around 1 cm ventral to infralimbic cortex layers V/VI, slightly dorsal to the nucleus accumbens core, which is the location these fibres have been reported to run through just prior to innervating the mPFC (52). The stimulating electrode was made by twisting together two wires made of Teflon-insulated stainless steel, approximately 125 µm in diameter with bare ends (Advent RM, UK). Glass recording electrodes were pulled to a tip diameter of 10-15 µm with an impedance c.1 MΩ. Two recording electrodes were filled with aCSF solution and individually placed in layers II/III and V of the infralimbic cortex. These were the recording areas of choice due to the relative preservation of hippocampal fibre connectivity in the slice preparation compared to the more dorsal prelimbic cortex (52).

*Extracellular recordings.* Local field potentials in response to constant 10-100 V stimulation were recorded from layers II/III and layer V, followed by paired-pulse responses to 100V stimulation with intervals of 20, 50, 100, 200 and 500ms. The maximal 100V stimulation was used to ensure optimal activation of local mPFC circuitry, enabling clear observation of any differences in response amplitudes between 3xTgs and controls. Five paired-pulse response traces were obtained per interval. Traces were subsequently averaged, and amplitudes and latencies of responses measured using Signal software (version 5; CED, UK).

### Bulk RNA sequencing

*Tissue extraction.* Bulk RNA sequencing analysis was performed on mPFC tissue from 3– and 6-month-old 3xTg and control mice (n= 4 per age, per group). Mice were first euthanised by cervical dislocation. Brains were obtained and sliced in RNAase-free PBS using a microtome (Leica VT1200), to obtain 1-2 slices containing the prefrontal cortex (AP 1.3 to 2.0). Sections of mPFC were punched out using thin-walled borosilicate glass capillary tubes (length of 75 mm, outer diameter of 1.5 mm, inner diameter of 1.1 mm; World Precision Instruments). Once extracted, tissue samples were placed inside labelled Eppendorf tubes which were kept at 4°C prior to RNA extraction.

*RNA extraction.* To extract RNA, tissue cores were first mixed with Invitrogen™ TRIzol™ Reagent (Invitrogen, Catalog No. 15596018). They were then mechanically dissociated using a 6 mm insulin syringe, and vortexed until cells appeared homogenized. Cells were centrifuged for 15 minutes at 15,000g at 4°C and the aqueous layer containing RNA was extracted. Pure isopropanol was added to purify the RNA in the precipitate. This was followed by spinning down for 10 minutes at 16,000g at 4°C. The supernatant was then removed from the pellet and 70% ethanol was used to wash the pellet twice. Lastly, the pellet was dissolved using ultrapure water. The resulting RNA solution was frozen and transported by dry ice.

*Library preparation and analysis.* Library preparation and sequencing was performed by NovogeneAIT (Singapore). Briefly, sequence reads were trimmed and aligned with HISAT2 (version 2.0.5). Genes were mapped to Mus musculus genome (GRCmm39) to generate uniquely mapped read counts. Read counts were converted to fragments per kilobase of transcript per million (FPKM) mapped reads values, and subsequently fed into DeSeq2 (version 1.20.0) to generate FoldChange (FC) values (120). Functional analyses were performed using the clusterProfiler library (version 3.8.1) (121). All graphs were plotted in RStudio (version 2023.09.1 +494).

### Statistics

4CGT and electrophysiology data were first tested for normality using the Shapiro-Wilk test, and equal variance using the F test. Unless stated, normally distributed 4CGT and electrophysiological data were analysed using a mixed-effects analysis or 2-way ANOVA respectively, with multiple comparisons (Bonferroni post hoc analysis). All testing was performed with Prism software (version 9.5.1, GraphPad UK). For behavioural data, if a mouse completed less than 10 trials over a session, this individual performance was excluded from the grouped analysis. Control and 3xTg performance were only analysed for n= 4 animals or more per group (i.e., 4 or more animals had performed 10 trials or more over the session).

## Acknowledgements

The authors would like to thank Mr Matthew Burgess for his expertise in helping set up behavioural testing chambers. This study was supported by A*STAR under it’s A*STAR Research Attachment Programme (ARAP). S.J. is supported by the A*STAR JCO-VIP award (Joint Council Office Grant BMSI/15-800003-SBIC-OOE).

## Author contributions

J.G. conceptualised the project; G.C., J.T., and J.G. designed the experiments; G.C. performed experiments and analyses relating to behavioural and electrophysiological data; G.C. and L.Y.T. performed RNA extraction and analyses relating to RNA sequencing data; G.C., J.G., and J.T. interpreted the results; G.C. wrote the original draft of the manuscript; all authors helped critically review the manuscript; J.G., J.T., and S.J. provided supervision over the project; J.G. and S.Y. facilitated the acquisition of funding.

## Data Availability

The datasets generated during and/or analysed during the current study are available from the corresponding author on reasonable request.

